# Adenylosuccinate Lyase activity in the Purine recycling pathway is essential for developmental timing, germline maintenance and muscle integrity in *C. elegans*

**DOI:** 10.1101/277640

**Authors:** Roxane Marsac, Benoît Pinson, Christelle Saint-Marc, María Olmedo, Marta ArtalSanz, Bertrand Daignan-Fornier, José-Eduardo Gomes

## Abstract

Purine homeostasis is ensured through a metabolic network widely conserved from prokaryotes to humans. Purines can either be synthesized *de novo*, reused, or produced by interconversion of extant metabolites using the so-called recycling pathway. Although thoroughly characterized in microorganisms, such as yeast or bacteria, little is known about the regulation of this biosynthesis network in metazoans. In humans, several diseases are linked to purine biosynthesis deficiencies through yet poorly understood etiologies. Particularly, the deficiency in Adenylosuccinate Lyase (ADSL), one enzyme involved both in the purine *de novo* and recycling pathways, causes severe muscular and neuronal symptoms. In order to address the mechanisms underlying this deficiency, we established *Caenorhabditis elegans* as a metazoan model organism to study purine metabolism, while focusing on ADSL. We show that the purine biosynthesis network is functionally conserved in *C. elegans*. Moreover, ADSL is required for developmental timing and germline stem cell maintenance, and muscle integrity. Our results allow to ascribe developmental and tissue specific phenotypes to separable steps of the purine metabolic network in an animal model. Particularly, the muscle, germline and developmental defects are linked specifically to the ADSL role in the purine recycling pathway.

## Introduction

Purine biosynthesis pathways ensure the production and homeostasis of AMP and GMP in the cell, and are widely conserved throughout evolution. The *de novo* pathway leads to the synthesis of IMP (Inosine monophosphate) using PRPP (5-Phosphoribosyl-1- pyrophosphate) as precursor. The recycling pathway (*a.k.a.* salvage pathway) can then transform extant purines, including IMP synthesized through the *de novo* pathway, to produce AMP and GMP (Figure 1A, reviewed in [1]). Importantly, if available in the extracellular environment, purines can be recovered by specific transporters and subsequently be metabolized through the recycling pathway, which requires much less energy expenditure. Indeed, purine synthesis through the *de novo* pathway requires the hydrolysis of six more molecules of ATP, than purine synthesis through the recycling pathway.

**Figure 1.**
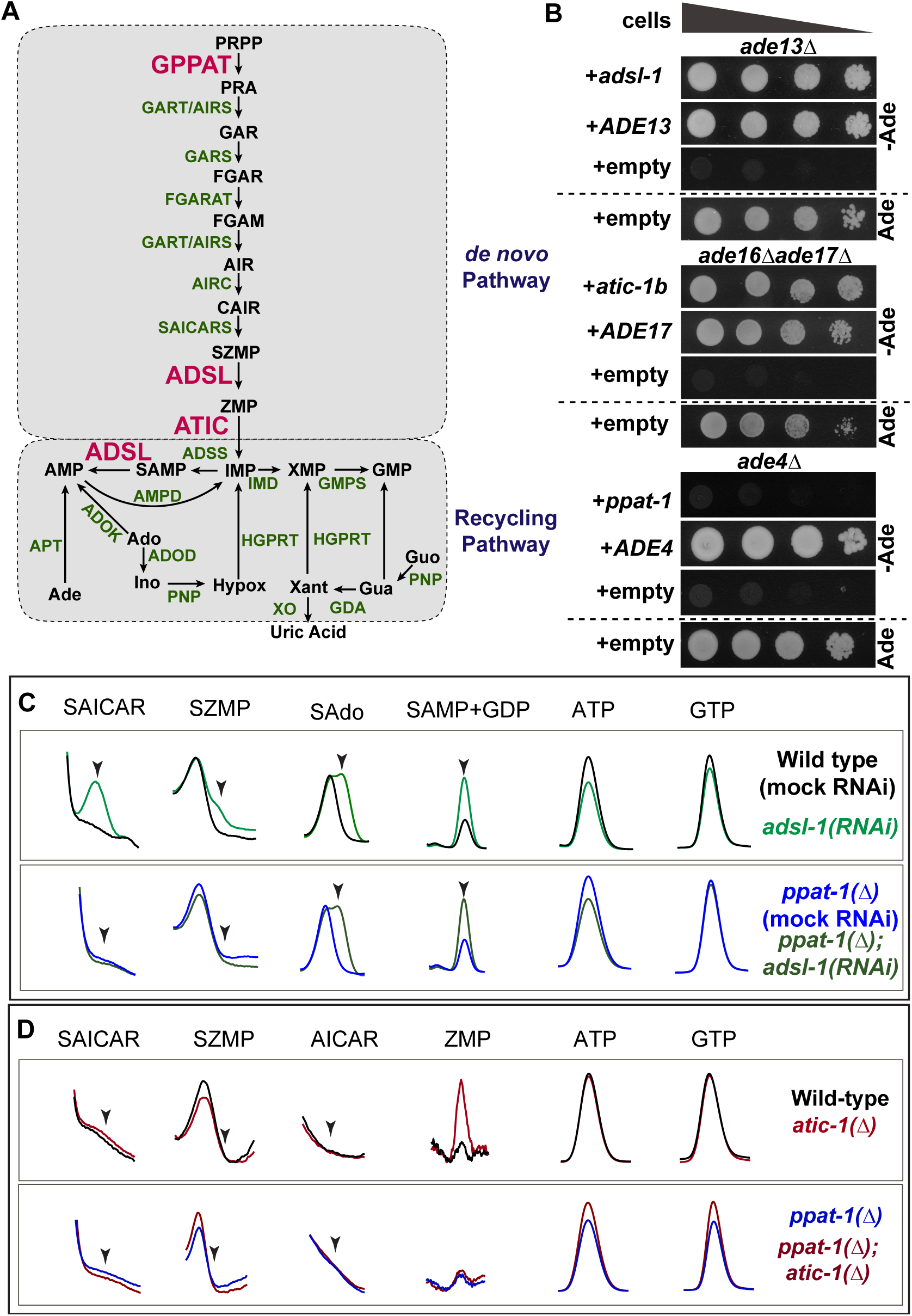
Purine metabolism in *C. elegans*. (A) Schematics of the Purine Biosynthesis pathways in *C. elegans* based on sequence homology. Enzymes subjected to functional analysis are represented in red, other enzymes in green and metabolites in black. (B) Drop test to assess adenine auxotrophy in yeast *S. cerevisiae* mutants deficient for enzymes ADSL (*ade13∆*), ATIC (*ade16∆ ade17∆*) or GPPAT (*ade4∆*) expressing respectively *C. elegans* genes *adsl-1*, *atic-1b* and *ppat-1*. Expression of *S. cerevisiae* genes *ADE13*, *ADE17* and *ADE4* was used as positive control, and expression vector without insert as negative control. A culture with adenine is shown as auxotrophy control. In all drop tests presented, four drops are shown per condition, corresponding to serial dilutions (1:10) of cellular suspensions, with deceasing cell concentrations from left to right. (C) Zoom in on HPLC chromatogram peaks of specific metabolites SAICAR (SZMP riboside form), SZMP, sAdo (succinyl-Adenosine, SAMP riboside form), SAMP, ATP and GTP, upon *adsl-1(RNAi)* and *ppat-1(∆); adsl-1(RNAi)*. (D) Zoom in on HPLC chromatogram peaks of specific metabolites SAICAR (SZMP riboside form), SZMP, AICAR (ZMP riboside form), ZMP, ATP and GTP, in *atic-1(∆)* and *ppat-1(∆); atic-1(∆)*. Abreviations: Ade - adenine, Ado - adenonsine, SAdo - succinyl-adenosine, Ino - Inosine, Hypox - hypoxantine, Xant - Xantine, Gua - guanine, Guo - guanosine. In panels (C) and (D), scales are adjusted differently among metabolites, in order to highlight differences between genotypes.

Imbalance in purine metabolism is involved in several diseases that have been extensively characterized, most of them associated with mutations in genes encoding for purine biosynthesis enzymes (reviewed in [2, 3]). The genetics and biochemistry of the purine biosynthesis are well established in unicellular organisms like bacteria or yeast, and are highly conserved. Nonetheless, how the disease-causing enzymatic deficiencies lead to the symptoms observed in patients remains elusive. It is particularly challenging to untangle the contribution of shortage in the final products - AMP and GMP - from mutation-caused accumulation of intermediate metabolites in the pathway. For instance, ZMP (5- amino-4-imidazole carboxamide ribonucleotide monophosphate [4]) and SZMP (succinyl-ZMP), two intermediates of the purine *de novo* pathway (Figure 1), are known to act as signal metabolites regulating gene expression transcriptionally in yeast [5]. Moreover, ZMP is known to activate AMPK (AMP activated kinase) by mimicking AMP [6] and SZMP has been shown to directly bind and interfere with pyruvate kinase activity in cancer cells [7]. In addition, SAMP (succinyl-AMP), an intermediate metabolite in the recycling pathway, has been shown to stimulate Insulin secretion in pancreatic cells [8]. Hence SZMP, ZMP or SAMP buildup could potentially disrupt homeostasis upon deficiencies in the enzymes using them as substrates - ADSL (SZMP and SAMP) and ATIC (ZMP, Figure 1A). Accordingly, mutations in the human genes encoding for these two enzymes are associated with ADSL deficiency and ATIC deficiency (*a.k.a.* AICAribosiduria), syndromes whose etiology remains unknown [9–14]. Adding to the complexity of the issue, ADSL catalyzes two different reactions, one in the *de novo* and one in the recycling pathway (Figure 1A), raising the question of which of the pathways is associated with the ADSL deficiency phenotypes.

The study of purine metabolic deficiencies in patients, faces yet other challenges, in comparison with microorganisms. One such challenge is tissue specificity, as different tissues may require different levels of purines, depending on their specialized function. For instance, in ADSL-deficiency patients, the most severe symptoms affect the muscles and the nervous system, suggesting these tissues have a larger requirement for purine metabolism, either for their function or during development. Another challenge is purine transport across tissues; the tissue affected by the deficiency not being necessarily the tissue where the enzyme is active. Moreover, purines can be available exogenously from the food source, therefore the consequences of purine metabolism deficiencies are not necessarily caused by limiting amounts of purines. Finally, purine metabolism might be subject to different regulations throughout development, in which case the impairments observed could be the consequence of a requirement for purine regulation at a specific developmental stage.

In order to address these issues, we sought to take advantage of *C. elegans* as model organism. Through a genetic approach, we characterized deficiencies in the purine biosynthesis pathways, focusing our analysis on an ADSL deficient mutant. Our goal was to establish the phenotypes associated with this enzymatic deficiency in order to uncover what functions, tissues and developmental stages are affected. Given ADSL is required both in the *de novo* and recycling pathways, we performed a comparative analysis with ATIC and GPPAT deficiencies to dissect the contribution of the *de novo* vs. recycling pathways to the phenotypes observed. We established that ADSL deficiency strongly disrupts muscle integrity, germline stem cells (GSCs) maintenance and post-embryonic development through the recycling pathway. By contrast, we found no essential requirement for the *de novo* pathway in *C. elegans* under standard laboratory conditions.

## Results

### Purine biosynthesis pathway is functionally conserved in *C. elegans*

In order to establish *C. elegans* as a model to study purine metabolism, we first sought to investigate the conservation of the purine biosynthesis pathways. We identified, through sequence homology, genes encoding the putative nematode enzymes involved in purine biosynthesis (Supplementary Figure 1A). We found that both *de novo* and recycling pathways are conserved in *C. elegans*, judged by sequence similarity. Specifically, for each enzyme we identified one single orthologue, in all but one case (three genes putatively encoding Xanthine Oxidase). Noteworthy, in *S. cerevisiae* the histidine pathway contributes to the purine *de novo* pathway, ZMP being a byproduct of histidine synthesis (reviewed in [1]). We have found no *C. elegans* orthologues for yeast HIS1, HIS4, HIS6 and HIS7 - required for ZMP synthesis through the histidine pathway - consistent with the histidine pathway loss in worms previously reported [15], thus ZMP is most likely synthesized only through the purine *de novo* pathway.

Given the sequence conservation, we undertook a functional analysis focusing on the ADSL enzyme, and including ATIC and GPPAT for comparison (see below). The genes encoding ADSL, ATIC and GPPAT in *C. elegans* were named respectively *adsl-1, atic-1* and *ppat-1* (<u>wormbase.org</u>). We first tested whether those three *C. elegans* enzymes were functionally conserved using an heterologous rescue assay. We transformed yeast *Saccharomyces cerevisiae ade13∆*, *ade16∆ ade17∆* and *ade4∆* mutants with plasmids containing the wild-type sequences of the predicted *C. elegans* homologous genes: *adsl-1*, *atic-1* and *ppat-1*, respectively. Expression of *adsl-1* restored adenine prototrophy of yeast *ade13∆* (Figure 1B), likewise expression of splicing isoform *atic-1b* restored adenine prototrophy of *ade16∆ade17∆* double mutant (Figure 1B). These results confirm *adsl-1* encodes the ADSL enzyme and *atic-1* encodes for the ATIC enzyme. By contrast, expression of *ppat-1* alone was not sufficient to restore adenine prototrophy of yeast *ade4∆* (Figure 1B), thus in our heterologous rescue assay we could neither confirm nor rule out *ppat-1* encodes the enzyme GPPAT.

We next tested functional conservation of ADSL, ATIC and GPPAT activities *in vivo*. For our functional analysis, we used deletion mutant alleles henceforth referred to as *adsl-1(∆)*, *atic-1(∆)* and *ppat-1(∆)* (see Materials and Methods). One would predict accumulation of ADSL substrates SZMP and SAMP upon *adsl-1* loss-of-function as well as ZMP buildup in *atic-1(∆)*, and no detectable *de novo* pathway intermediate metabolites in *ppat-1(∆)* (Supplementary Figure 1B). To test these hypotheses, we performed worm metabolic profiling by HPLC (Supplementary Figure 1C). Importantly, sequence alignment indicates ADSL, ATIC and GPPAT are highly conserved (Supplementary Figure 2). We observe in our metabolic profiling a consistent, accumulation of SZMP, and its riboside form SAICAR, upon *adsl-1(RNAi),* as well as a robust accumulation of SAMP, and its riboside form SAdo (Figure 1C), while neither metabolite was detected in N2 reference strain (henceforth wild-type), demonstrating that both *de novo* and recycling pathways are functionally conserved in *C. elegans*, and *adsl-1* is required for ADSL activity *in vivo*. Furthermore, we observe a modest but consistent accumulation of ZMP both in *atic-1(*∆) mutants (Figure 1D), as well as *atic-1(RNAi)* animals (not shown), demonstrating *atic-1* encodes ATIC enzyme. The accumulation of SZMP and ZMP in *adsl-1(RNAi)* and *atic-1(∆)* was rather modest, by contrast we observed a robust accumulation of SAMP in *adsl-1(RNAi).* These results suggests that *C. elegans* purine biosynthesis relies mostly on the recycling pathway, at least in standard culture conditions. Noteworthy, ATP levels are slightly lower in *adsl-1(RNAi)* compare to wild-type (Figure 1C), otherwise the abundance of ATP and GTP were not severely affected in the deficiencies analyzed.

**Figure 2.**
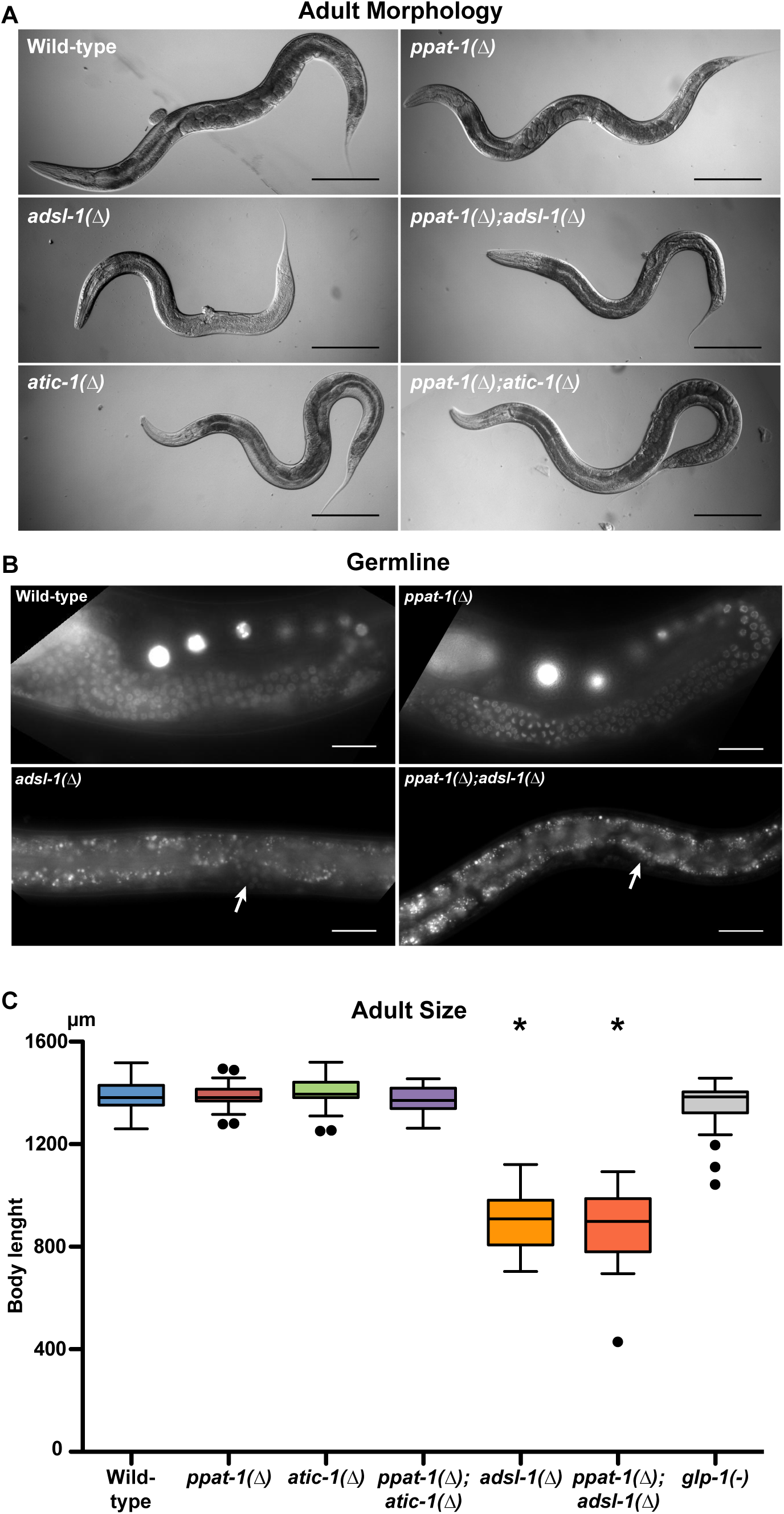
Developmental defects in purine mutants. (A) DIC whole body images of representative young adults for each of the genotypes studied (bar= 200 µm). (B) Fluorescence micrographs of worms expressing H2B::GFP in the germline under the control of the *pie-1* promoter. Arrows point at rare germ nuclei expressing H2B::GFP; spotty pattern throughout *adsl-1(∆)* and *ppat-1(∆); adsl-1(∆)* corresponds to intestinal auto-fluorescence (bar= 25 µm). (C) Tukey box plot depicting body length of all purine mutants and *glp-1(-)* for comparison (* - statistically significant difference relative to wild-type - one way ANOVA; n= 30 for all genotypes).

As to the conservation of GPPAT, the HPLC analysis provided crucial functional evidence: neither SZMP was detected in *ppat-1(∆); adsl-1(RNAi)* (Figure 1C) nor was ZMP detected in *ppat-1(∆); atic-1(∆)* (Figure 1D), or *ppat-1(∆); atic-1(RNAi)* (not shown). This demonstrates that *ppat-1* gene is indeed required for purine synthesis through the *de novo* pathway. Noteworthy, SAMP accumulation in *ppat-1(∆); adsl-1(RNAi)* is similar to *adsl-1(RNAi)* alone, providing an internal control for RNAi efficiency and further confirming SZMP accumulation in *adsl-1(RNAi)* is dependent on *ppat-1*. In summary, our metabolic profiling validates functional conservation of ADSL and ATIC, and demonstrates *ppat-1* is required for the *de novo* pathway, consistent with encoding the GPPAT enzyme in *C. elegans*.

### Functional analysis of *adsl-1*

Because SZMP and SAMP are known to function as regulatory metabolites in other organisms [5, 8, 16, 17] and because they may play an important role in the etiology of the ADSL deficiency - the enzyme metabolizing these two compounds, we focused our functional analysis on *adsl-1*. In order to distinguish the effects of SZMP buildup from those due to the loss of *de novo* purine synthesis, we further include *ppat-1* in our analysis. Moreover, in order to distinguish the effect of ADSL activity in the *de novo* pathway from the activity in the salvage pathway we used for comparison ATIC deficient worms. ATIC participates only in the *de novo* pathway and accumulates ZMP, which shares some targets with SZMP at least in yeast [5, 16]. Finally, in order to test whether phenotypes observed in *adsl-1(∆)* were due to SZMP buildup, we performed systematically epistasis analysis in the *ppat-1(∆); adsl-1(∆)* double mutant, and similarly to test whether phenotypes in the *atic-1(∆)* mutant were caused by ZMP accumulation we performed epistasis in the *ppat-1(∆); atic-1(∆)* double mutant. Throughout the manuscript, we refer collectively as “purine mutants” to all three single mutant homozygous animals, plus the *ppat-1(∆); adsl-1(∆)* and *ppat-1(∆); atic-1(∆)* double mutants; and as “*de novo*” pathway mutants to *ppat-1(∆)*, *atic-1(∆)* single mutants and *ppat-1(∆); atic-1(∆)* double homozygous mutant animals.

### adsl-1 is required for germline stem cells’ (GSCs) maintenance

The *adsl-1(∆)* homozygous animals display striking morphological defects, they are noticeably smaller than wild-type, sterile and present a protruding vulva (Figure 2A). We observe a complete absence of oocytes and sperm cells in *adsl-1(∆)* and *ppat-1(∆); adsl-1(∆)* worms, only a reduced number of nuclei morphologically resembling the GSCs are detectable near the vulva, where the germline distal tip cell is found in wild-type animals (Supplementary Figure 3A). By contrast, the homozygous mutants in the *de novo* pathway are viable and fertile, and the overall morphology is similar to wild-type controls. To better characterize the GSCs phenotype, we used a reporter transgene expressing H2B::GFP fusion in the GSCs under the control of the *pie-1* promoter [18]. While in *ppat-1(∆)* single mutants the germline is indistinguishable from wild-type, in *adsl-1(∆)* and *ppat-1(∆); adsl-1(∆)* animals sterility is fully penetrant and the germline is barely detectable if at all (Figure 2B). Indeed we observed 52% of *adsl-1(∆)* (n = 23) and *ppat-1(∆)*; *adsl-1(∆)* animals (n = 25) had no detectable H2B::GFP, 44% had 10 or less H2B::GFP positive nuclei, and 4% had barely above 10 H2B::GFP positive nuclei. Hence *adsl-1* is required for proliferation and/or maintenance of the GSCs, independently of *ppat-1*.

**Figure 3.**
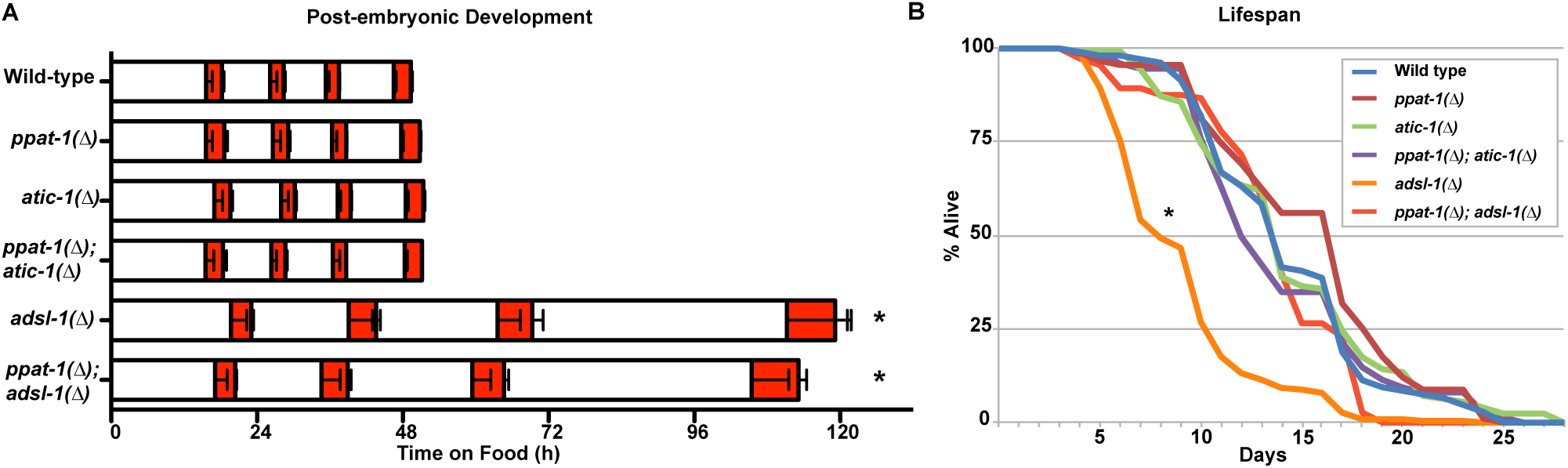
Life traits in purine mutants. (A) - Graph depicting duration of post-embryonic developmental stages, white bars represent larval stages, from L1 to L4, orange bars represent lethargus phases consecutive to each larval stage (* - statistically significant difference relative to wild-type - one way ANOVA; see detailed representation of the data in Supplementary Figure 4). (B) Graph depicting fraction of alive worms over time for all genotypes studied (* - statistically significant difference relative to wild-type - Kaplan-Meier; total combined of three independent experiments for each genotype, wild-type n= 106, *ppat-1(∆)* n= 91, *atic-1(∆)* n= 126, *ppat-1(∆); atic-1(∆)* n= 149, *adsl-1(∆)* n= 227, *ppat-1(∆); adsl-1(∆)* n= 113).

It has been shown that the sterility associated with a deficiency in the pyrimidine recycling pathway could be rescued by supplementation of the culture medium with pyrimidine nucleosides [19], we thus tested whether supplementation with purine nucleosides could rescue *adsl-1(∆)* sterility. When cultured with 20 mM Adenosine (Ado in Figure 1A) in the medium from hatching until adulthood, both *adsl-1(∆)* (n = 55) and *ppat-1(∆); adsl-1(∆)* (n = 54) were sterile, much like the non-supplemented controls (n = 80 and n = 88, respectively). As expected, 100 mM Inosine (Ino in Figure 1A) supplementation had no effect in *adsl-1(∆)* (n = 28) nor in *ppat-1; adsl-1(∆)* (n = 64) sterility. Under the same conditions wild-type (20 mM Adenosine n = 86; 100 mM Inosine n = 99) and *ppat-1(∆)* (20 mM Adenosine n = 76; 100 mM Inosine n = 100) worms were fertile, to the same extent as non-treated controls (wild-type n = 116; *ppat-1(∆)* n = 114). We found no evidence that purine supplementation in the culture medium allows to recover the GSCs defects of *adsl-1(∆)*.

As mentioned above, the *de novo* pathway mutants present a germline similar to wild-type, in contrast with the complete sterility of *adsl-1(∆)*. In *C. elegans*, adult hermaphrodites reproduce through self-fertilization, and the number of progeny being modulated by food source and metabolic status [20, 21]. In order investigated whether the *de novo* pathway could impact the level of fertility, we measured total number of eggs laid by *ppat-1(∆)*, *atic-1(∆)* and *ppat-1(∆); atic-1(∆)* hermaphrodites throughout their reproductive life. We observed that *ppat-1(∆)* fertility is in the wild-type range, while *atic-1(∆)* fertility is significantly lower (Supplementary Figure 3B). Importantly, *ppat-1(∆)* is epistatic to *atic-1(∆)* regarding fertility, suggesting that ZMP accumulation, rather than *de novo* pathway impairment, might reduce fertility. To test this hypothesis we cultured *ppat-1(∆); atic-1(∆)* in presence of ZMP precursor AICAR (0.1 mM) in the culture medium - under this condition internal ZMP builds up (see below). We observed a reduced fertility in *ppat-1(∆); atic-1(∆)* treated with AICAR, compared to the control (Supplementary Figure 3B), consistent with our hypothesis. Furthermore, wild-type worms cultured with 0.5 mM AICAR also displayed a reduced fertility. We conclude that ZMP buildup in *atic-1(∆)* causes reduced fertility, while the *de novo* pathway *per se* has no impact on this phenotype, as *ppat-1(∆)* is similar to wild-type. Taken together these data show that *adsl-1* role in the recycling pathway, and not in the *de novo* pathway, is essential for germline integrity.

The strong germline defects in *adsl-1(∆)* prompted us to investigate whether *adsl-1* is required during early embryonic development. We therefore examined the viability of ADSL deficient embryos. Given *adsl-1(∆)* are sterile, we analyzed the progeny of *adsl-1(∆)/+* heterozygotes; if *adsl-1* zygotic expression were essential in the embryo one would expect ~25% embryonic lethality, we observed only 7.7%. Therefore, either *adsl-1* is not essential for embryo development or maternal contribution is, at least in part, sufficient. We used RNAi to discriminate between these two hypotheses. Because *adsl-1* full penetrant loss-of-function leads to sterility, we performed *adsl-1(RNAi)* starting at the L4 stage resulting in a hypomorphic phenotype. Under these conditions we observed 32.2% embryonic lethality in *adsl-1(RNAi)* (Supplementary Figure 3C), hence maternal expression ensures *adsl-1* function during embryonic development. To test whether the *de novo* pathway contributed to embryonic lethality we performed *adsl-1(RNAi)* in the *ppat-1(∆)*, resulting in 31.5% embryonic lethality, similarly to *adsl-1(RNAi)* alone. The *de novo* pathway mutants, on the other hand, display normal embryonic development, just as in WT controls: nearly 100% of the laid eggs hatched in all the *de novo* pathway mutants,showing neither *ppat-1* nor *atic-1* are essential for embryo development. Thus *adsl-1* function in the recycling pathway is required for proper embryonic development.

### adsl-1 is required for body size

The overall morphology observed in *adsl-1(∆)* and *ppat-1(∆); adsl-1(∆)* (morphologically indistinguishable from *adsl-1(∆)* single mutants) suggests that cell lineage specification and morphogenesis are not affected, given somatic tissues are properly formed. No defects are observed in the pharynx, intestine or hypodermis (other than the vulva) of *adsl-1(∆)* and animals are viable and mobile. Hence the smaller size is not the result of defects in fate specification. To better understand this phenotype, we measured body length in adult worms. *adsl-1(∆)* are significantly smaller than wild-type, while the *de novo* mutants are indistinguishable (Figure 2C). *ppat-1(∆); adsl-1(∆)* are similar to *adsl-1(∆)*, thus ADSL-1 activity in the recycling pathway is responsible for the *adsl-1(∆)* body size phenotype. Given the volume occupied by the germline in adult *C. elegans* we wondered whether sterility could account for the small size in *adsl-1(∆)*, we therefore compared *adsl-1(∆)* to *glp-1(e2141)* mutants, known to lack a germline when cultured at 25°C [22]. Given adult *glp-1(e2141)* length is in the wild-type range (Figure 2C), sterility cannot account for the small size of *adsl-1(∆)*. Noteworthy, newly hatched *adsl-1(∆)* animals are the same size as wild-type and morphologically indistinguishable (not shown), hence size defects manifest during post-embryonic development. We conclude *adsl-1* is required, through the recycling pathway, for adult body size independently of the sterility phenotype, presumably by affecting cell size (see discussion).

### - adsl-1 is required for developmental timing, but not lifespan

The small size of *adsl-1(∆)* prompted us to investigate whether post-embryonic development was impaired. Given newly hatched *adsl-1(∆)* are indistinguishable from wild-type, we hypothesized the growth defects were linked to abnormal larval development. Upon hatching and before adulthood, *C. elegans* goes through four larval stages each followed by an inactive state known as lethargus during which a molt takes place [23]. Importantly, this sequence follows a highly reproducible timing under standard culture conditions [24]. We therefore measured timing of post-embryonic development, in purine mutants, using the method described by Olmedo *et al* [25]. Briefly, newly hatched L1 larvae carrying a transgenic luciferase are cultured in presence of luciferin substrate, emitting luminescence when active [26]. Since larvae cease feeding during lethargus, they stop taking the substrate and a drastic drop in luminescence is observed. Hence, monitoring the luminescence of individual larvae allows to precisely time the occurrence of each larval (high luminescence) and lethargus (low luminescence) phases (Supplementary Figure 4A; [25]). We observed that both *adsl-1(∆)* and *ppat-1(∆); adsl-1(∆)* present a strongly delayed post-embryonic development (Figure 3A, Supplemental Figure 4B). Although all larval and lethargus stages are affected in *adsl-1(∆)* mutants, the delay is more pronounced as post-embryonic development progresses. By contrast, the overall timing of post-embryonic development is not affected in *de novo* pathway mutants. We therefore conclude that *adsl-1* function in the recycling pathway is required for normal post-embryonic development.

**Figure 4.**
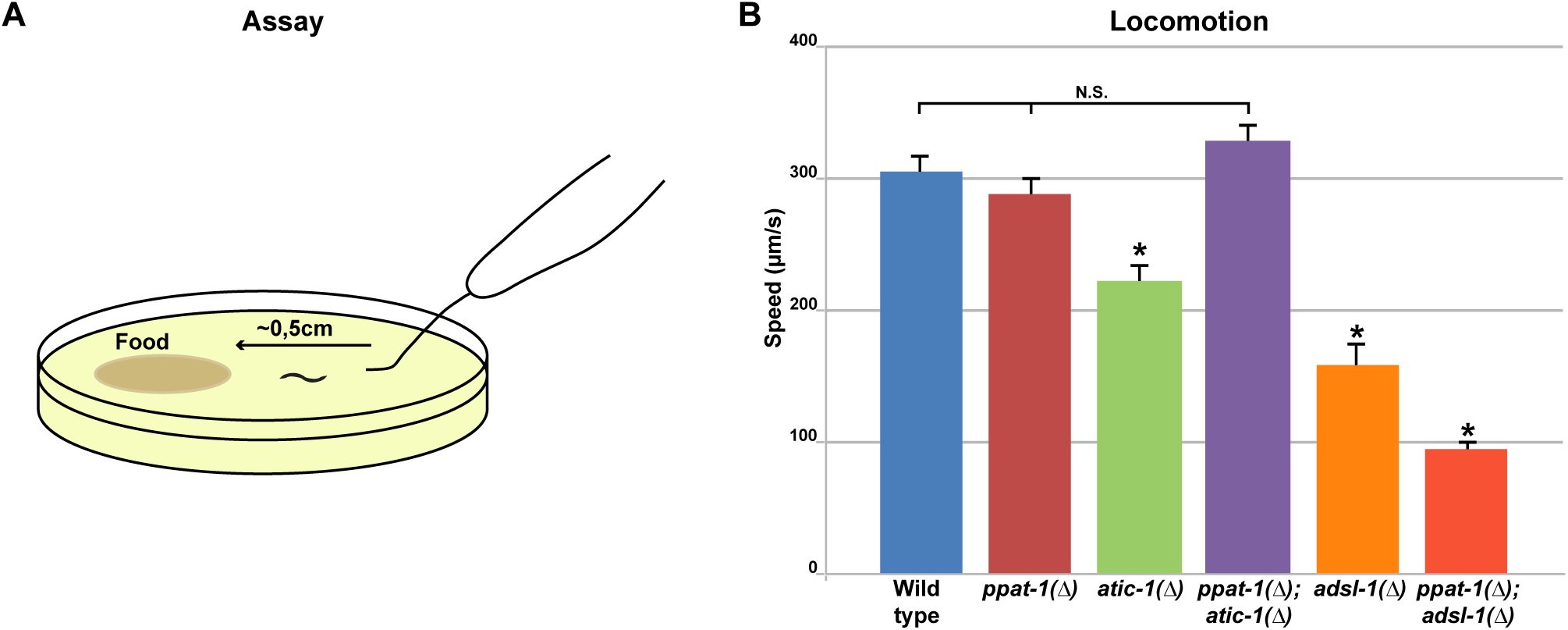
Locomotion of purine mutants. (A) Cartoon schematics of the locomotion behaviour assay. (B) Graph depicting the average locomotion speed in all purine mutants (error bar - standard error of the mean; n = 30 in all genotypes except wild-type and *ppat-1(∆); atic-1(∆)* n= 40; * - statistically significant difference relative to wild-type - Student’s t-test).

Post-embryonic development, metabolism and the germline are all known to impact longevity in *C. elegans* (reviewed in [27]). Because of *adsl-1(∆)* sterility and developmental delay we measured the lifespan of the purine mutants. *adsl-1(∆)* mutant has a shorter lifespan that wild-type (Figure 3B). By contrast, the *de novo* pathway mutants’ lifespan is in the wild-type range (Figure 3B). Importantly, *ppat-1(∆)* suppresses the *adsl-1(∆)* lifespan phenotype as *ppat-1(∆); adsl-1(∆)* double mutant survival profile is in the wild-type range (Figure 3B). These results strongly suggest that SZMP buildup in *adsl-1(∆)*, rather than ADSL activity in the recycling pathway, is responsible for the reduced lifespan. Moreover, we conclude that the developmental delay and sterility of *adsl-1(∆)* are not linked to lifespan.

### Imbalance in purine metabolism impacts locomotion

Given the severe muscular and neuronal symptoms of ADSL deficiency patients [9, 11, 28], we sought to investigate muscle and neural defects in *adsl-1(∆)* worms. In order to address muscle function, we performed a locomotion assay (Figure 4A; [29]), and observed that *adsl-1(∆)* mutants were slower than WT (Figure 4B). *atic-1(∆)* mutants, were also slower than wild-type, albeit to a lesser extent (Figure 4B), suggesting the *de novo* pathway could have a role in locomotion. However, *ppat-1(∆)* animals were indistinguishable from wild-type, thus *de novo* pathway *per se* does not affect locomotion. Moreover, the locomotion defect in *adsl-1(∆)* was not suppressed by *ppat-1(∆)*, *ppat-1(∆); adsl-1(∆)* was actually slower than *adsl-1(∆)* single mutant. Thus, ADSL activity in the recycling pathway, rather than SZMP buildup, accounts for the *adsl-1(∆)* locomotion defect. By contrast, *atic-1(∆)* locomotion defect was suppressed by *ppat-1(∆)* as *ppat-1(∆); atic-1(∆)* speed was in the wild-type range. Hence, slower locomotion in *atic-1(∆)* is linked to ZMP build up, rather than the *de novo* pathway *per se*.

### AICAR treatment

The ADSL product ZMP is known to activate AMPK by mimicking AMP [6], and AICAR treatment is known to impact muscle function in mice, presumably through ZMP induced AMPK activation [30]. To test whether ZMP is responsible for the locomotion defects in *atic-1(∆)*, we tested whether providing the worms with external AICAR would impact locomotion. We treated late L4 stage wild-type and *de novo* pathway mutants with AICAR, for 24 hours. Somewhat surprisingly AICAR treatment did not significantly affect locomotion, neither among the genotypes tested, nor comparing with untreated control (Supplementary Figure 5A). Given this surprising result, we performed metabolic profile analysis of AICAR treated worms. As expected, we observed higher levels of AICAR upon treatment, and noticeably a significant buildup of both ZMP and ZTP in all treated genotypes (Supplementary Figure 5B), showing AICAR was efficiently internalized and phosphorylated. As expected ZMP levels are higher in *atic-1(∆)* than in other genotypes due to cumulative buildup of ZMP from the *de novo* pathway and from AICAR treatment (Supplementay Figure 5C). Therefore buildup of ZMP in adult worms *per se* does not explain the reduced locomotion observed *atic-1(∆)*. Importantly, while AICAR treatment did not significantly affect ATP levels across genotypes, it lead to an increase in uric acid, particularly in *atic-1(∆)* and *ppat-1(∆); atic-1(∆)*. Moreover, we also observed, in all treated genotypes, an accumulation of ADSL substrate SZMP and its riboside form SAICAR. Therefore, AICAR treatment leads to increased levels of ZMP as expected, but also of uric acid, ZTP and SZMP (Supplementary Figure 5B). The SAICAR and SZMP accumulation indicates that not only SZMP to ZMP conversion by ADSL is inhibited by AICAR treatment, this reaction can even be reversed, given SZMP was detected in *ppat-1(∆)* and *ppat-*1*(∆); atic-1(∆)* mutants (Supplementary Figure 5B, see Supplementary Figure 1B). Accordingly,in yeast ADSL enzyme has been shown to convert ZMP in SZMP when ZMP is in excess [31], which seems to also be the case for *C. elegans* ADSL.

**Figure 5.**
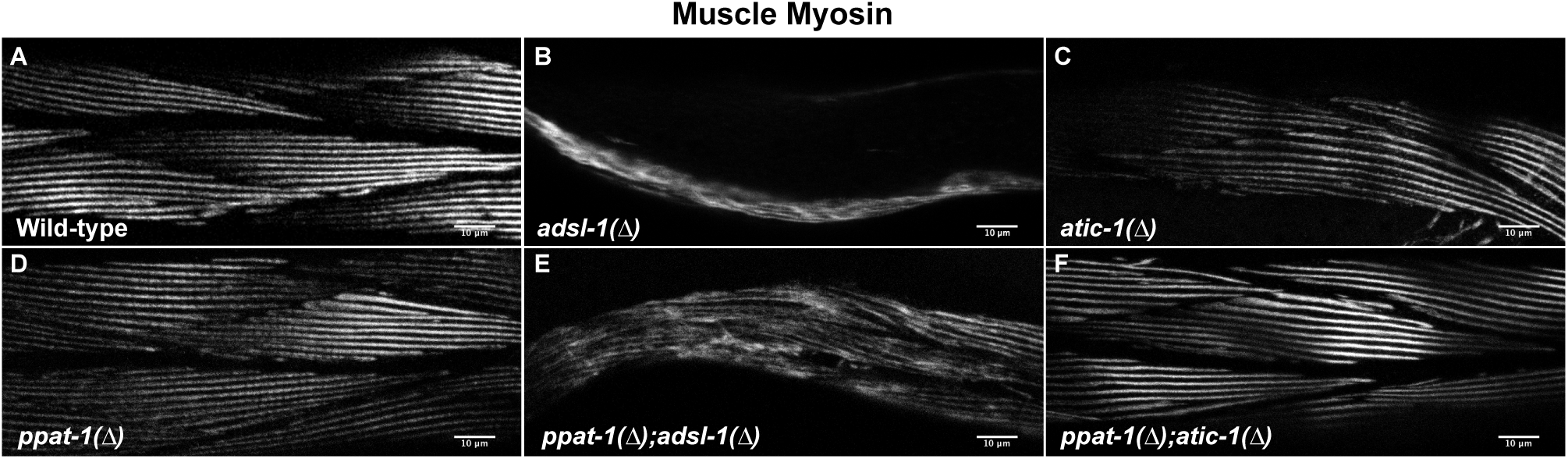
Muscle structure in purine mutants. Confocal micrographs of muscle myosin immunostaining in young adult’s body wall muscle cells of all studied purine mutants (bar= 10µm). (A) Wild-type. (B) *adsl-1(∆)*. (C) *atic-1(∆)*. (D) *ppat-1(∆)*. (E) *ppat-1(∆); adsl-1(∆)*. (D) *ppat-1(∆); atic-1(∆)*.

### Muscle structure, but not the neuromuscular junction, is disrupted in *adsl-1(∆)*

One possible explanation for the locomotion defects observed in *atic-1(∆)* and *adsl-1(∆)* is that the neuromuscular junction is compromised. We therefore tested whether purine mutants were sensitive to Levamisole and Aldicarb. Both drugs activate acetylcholine receptors at the neuromuscular junction in *C. elegans*, at either the pre-synapse (Aldicarb) or post-synapse (Levamisole) leading to paralysis of wild-type worms [32], while mutants with defective cholinergic neuromuscular junction are not affected [33, 34]. We found no significant differences among purine mutants and wild-type (Supplementary Figure 6A and B), demonstrating that the neuromuscular junction remains functional upon purine metabolism deficiencies. We therefore tested whether other neuronal functions might be impaired in purine mutants. We performed a standard touch assay as described in the literature [29]. We observed a similar response to touch in all purine mutants as in the wild-type, thus in our experiments the mechanosensory function as well as the behavioral response are intact in purine mutants (Supplementary Figure 6C). In summary we found no evidence for neuronal defects in the purine metabolism mutants tested.

**Figure 6.**
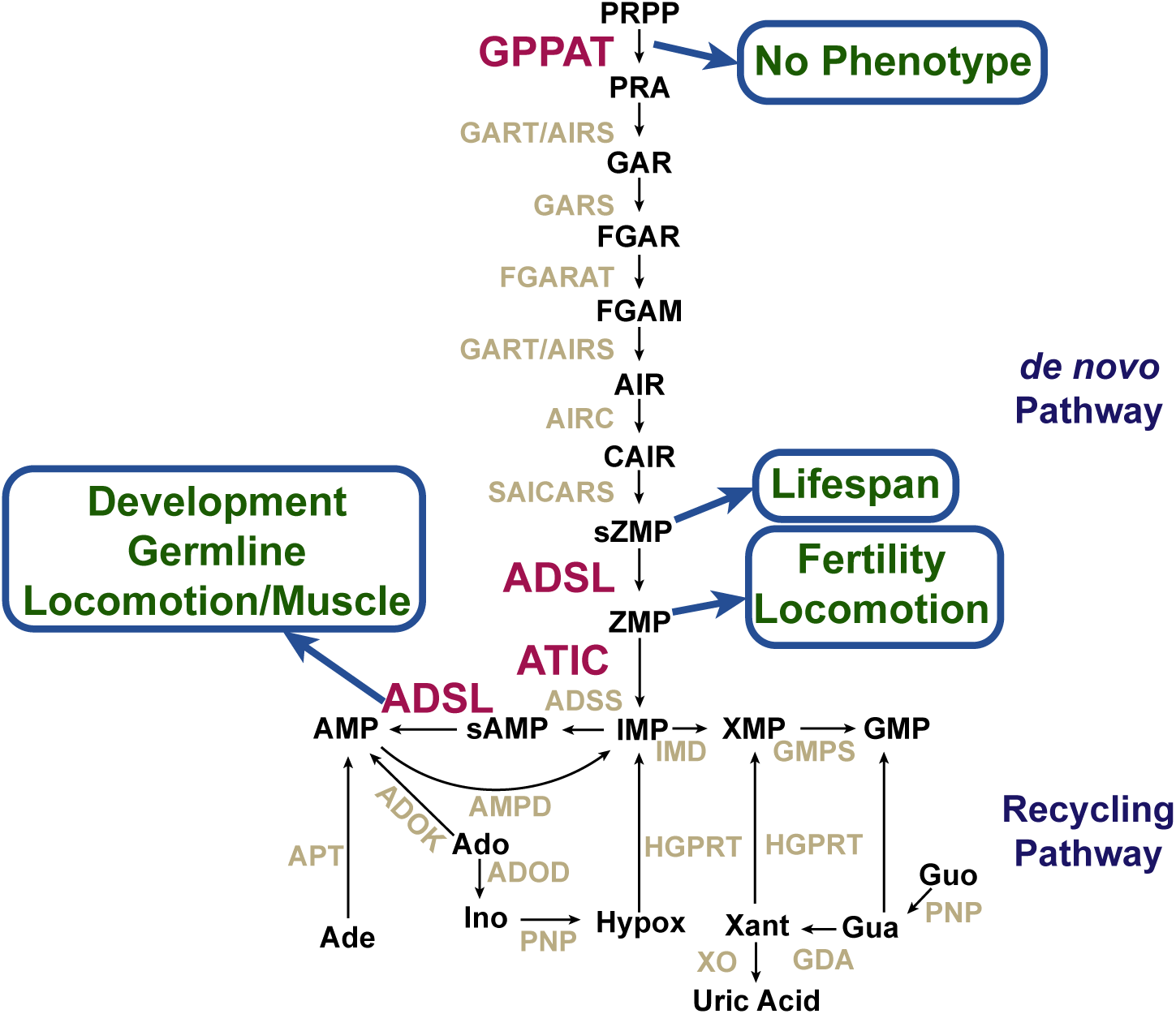
Summary of phenotypes linked to deficiencies in the purine biosynthesis pathway in *C. elegans* reported in this study.

Given these results, we hypothesized that the locomotion defects we observed could be explained by muscle malfunction. We performed an analysis of muscle cell structure through immunohistochemistry and confocal microscopy. As expected, we observed regular parallel arrangement of myosin fibers in the wild-type (Figure 5A), while in *adsl-1(∆)* mutants (n = 13), muscle myosin fibers formed an irregular meshwork and muscle mass seemed reduced (Figure 5B). In *atic-1(∆)* (n = 14), by contrast, the structure of muscle myosin fibers was indistinguishable from wild-type controls (Figure 5C, n = 10), as are *ppat-1(∆)* (Figure 5D, n = 13). Furthermore *ppat-1(∆); adsl-1(∆)* (Figure 5E, n = 10) displayed the same muscle fiber defects as *adsl-1(∆),* and *ppat-1(∆); atic-1(∆)* were indistinguishable from *ppat-1(∆)* and *atic-1(∆)* single mutants (Figure 5F, n = 15), showing that *de novo* pathway deficiency does not affect muscle structure (Figure 5). These results indicate that ADSL role in the recycling pathway is required for muscle integrity, likely underlying the locomotion defects we observed.

## Discussion

### A nematode model to study ADSL deficiency and more broadly purine metabolism

We describe here the phenotypes and metabolic profiles associated with ADSL deficiency in the nematode *C. elegans*. This study provides a reference to future studies investigating the underlying molecular mechanisms of the phenotypes observed. Although phenotypes are to some extent genetically separable, we show the most severe ones are linked to ADSL activity in the recycling pathway (Figure 6). This points to the conversion of S-AMP in AMP as the critical enzymatic step in ADSL deficiency. For instance, there is a long standing debate concerning the etiology of the pathology associated with ADSL deficiency. It has been proposed that the SAICAR/S-Ado ratio correlated with specific symptoms [14, 28], while more recent studies found no evidence for such correlation [10, 12]. Studies in human subjects are of limited reach to solve these types of questions, while a genetically amenable model organism shall allow to investigate molecular mechanisms underlying specific phenotypes, and potentially provide useful information to the understanding of the disease. *C. elegans* as metazoan model organism provides the means to investigate the metabolic regulation of developmental progression, muscle defects or cell proliferation in the context of ADSL deficiency, as well as in other deficiencies in the purine biosynthesis pathway.

### Purine recycling and GSCs maintenance

One striking phenotype of *adsl-1(∆)* is the severe reduction in GSCs, leading to sterility. In adult *C. elegans* the germline is the sole proliferating tissue. More precisely, GSCs in the distal gonad proliferate under the control of Notch signaling from neighboring somatic cells, while in the proximal region the germline nuclei enter meiotic differentiation (reviewed in [35, 36]. Interestingly, deficiencies in both the pyrimidine [19]) and pyridine nucleotides (*a.k.a.* NAD+) recycling pathways [37] have also been linked to germline defects in *C. elegans*, suggesting nucleotide metabolism to play a critical role in maintaining the GSCs pool. Importantly however, *adsl-1(∆)* phenotype is more severe than in either case. In *pnc-1(-)* mutants, affecting the pyridines salvage pathway, animals are fertile despite germline defects. In *cdd-1/2(-)* mutants, deficient for the Cytidine Deaminase in the pyrimidine recycling pathway, GSCs proliferation could be restored with pyrimidine supplementation. By contrast, purine supplementation had no effect on *adsl-1(∆)* GSCs. While one cannot exclude the defects in these mutants are unrelated, the three metabolic pathways are closely connected, it is therefore plausible that germline defects in *adsl-1(-)*, *pnc-1(-)* and *cdd-1/2(-)* share a common mechanism, through crosstalk among these metabolic pathways. In that case, the severity of the *adsl-1(∆)* phenotype remains to be explained. One tempting explanation is that the primary metabolic defect lies on the purine pathway, and that imbalance in the other nucleotide biosynthesis pathways impact purine metabolism as an indirect consequence. It is known that GSCs proliferation is influenced by environmental factors, particularly nutrient availability, and is likely regulated by metabolic signals (reviewed in [38]). Our results are consistent with the purine recycling pathway serving as sensor integrating nucleotide pathways’ status, and providing a metabolic signal regulating GSCs proliferation.

Interestingly, the purine biosynthesis pathway has been shown to play a role in the balance between proliferation and differentiation in both stem cells and cancer cells, with inhibition of purine synthesis leading to increased differentiation at the expense of proliferation [39, 40]. A similar mechanism could explain *adsl-1(∆)* GSCs defect, in that scenario the ADSL deficiency would lead to germline nuclei entering meiosis at the expense of proliferation.

### Purine metabolism and development

Metazoans often control their development in response to environmental conditions, namely nutriment availability. This control involves metabolic regulation, for instance through Insulin/Insulin-like or mTOR pathways (reviewed in [41]). In *C. elegans* the adult worm size varies depending on the bacterial food source [42], suggesting size is one of the developmental phenotypes subjected to metabolic regulation. This regulation could, in theory be due to metabolic regulation of cell fate decisions, or alternatively be a consequence of metabolic regulation of cell size. In *adsl-1(∆)*, based on our analysis, only GSCs and vulval cells present cell fate abnormalities, which have no impact in overall adult size. Hence the small size we observe is most likely due to an effect of purine metabolism on cell size.

In addition to size, *adsl-1(∆)* also displays a strong delay during post-embryonic development. In *C. elegans* progression through larval stage and molting cycles is highly regulated, involving oscillations of LIN-42, the orthologue the circadian protein PERIOD [43, 44], reviewed in [45]. The developmental changes associated with this progression involve a tight coordination of different cell lineages across the worm’s body, and are very well characterized, as are the genetic pathways involved (reviewed in [46]). By contrast, the upstream mechanisms and signals triggering or modulating these developmental transitions are poorly understood. The quality and amount of food available certainly play a role, as they have been shown to impact the timing of post-embryonic development [21, 47]. The dramatic developmental delay in *adsl-1(∆)* points at a metabolic regulation of these developmental transitions involving the purine recycling pathway.

One possibility could be that purine synthesis deficiency in *adsl-1(∆)* is limiting ATP levels, causing slow development and smaller size. Moreover, we have observed that in *adsl-1(∆)* food uptake is reduced compared to wild-type [48], thus not only ATP but limiting overall bulk nutrients could account for the phenotypes observed. Although limiting nutriments and ATP levels are an important factor, and certainly have a sizable impact, several elements suggest this hypothesis does not suffice to explain the *adsl-1(∆)* defects. In *eat* mutants, known to have feeding impairment, food uptake defects don’t always correlate with developmental delay (M.O. and M.A.S. unpublished results). Moreover, in *C. elegans* unfavorable environmental conditions, namely food shortage, induce an alternative larval stage known as dauer [23]. If food intake is limiting in *adsl-1(∆),* it is not so low as to mimic the nutrient scarcity causing dauer formation. Our observation that *adsl-1(∆)* progresses through post-embryonic development without entering the dauer stage is consistent with bulk nutriments not being the limiting factor. As to ATP levels, our metabolic profiles show them to be relatively high in *adsl-1(RNAi)*, only slightly reduced compared to wild-type. This is not so surprising given adenine or adenosine are available from the bacterial food source could be used to produce ATP despite ADSL deficiency (see Figure 1A). An alternative hypothesis is that post-embryonic developmental and adult size are regulated by the wiring of the purine recycling pathway, rather than the synthesis of its final products. In this view, purine intermediate metabolites could function as metabolic sensors modulating developmental progression.

### Purine metabolism and neuro-muscular function

Both neuronal and muscle tissues are severely affected in purine metabolism syndromes, suggesting these two tissues are particularly susceptible upon purine metabolism impairment in patients. In our nematode model, we observed locomotion defects associated with both ATIC and ADSL deficiencies. Somewhat surprisingly, no discernible neuronal damage was observed in any purine mutant. All of the genotypes tested displayed sensitivity to Levamisole and Aldicard, similarly to wild-type controls, showing neuromuscular junction was still functional. Moreover, mechanosensory response wasn’t affected either in purine mutants. Hence, we could not identify neuronal dysfunctions to account for the locomotion defects.

We did however find evidence for muscle abnormalities that could explain the locomotion defects. Noteworthy, the locomotion defects we observed in *atic-1(∆)* and *adsl-1(∆)* were linked to different mechanisms. Contrary to *adsl-1(∆)*, the locomotion delay in *atic-1(∆)* was suppressed in the *ppat-1(∆); atic-1(∆)* double mutant, hence linked to ZMP buildup. One possible explanation is that ZMP accumulation leads to metabolic changes that in turn impact muscle function, possibly through AMPK activation [6]. Importantly, the structure of muscle myosin fibers in *atic-1(∆)* are undistinguishable from the wild-type, thus muscle integrity seems not to be affected in *atic-1(∆)*. The ZMP effect on locomotion, however, is not straightforward; upon AICAR treatment of L4 larvae / young adults ZMP builds up as expected, nonetheless locomotion is not affected. Although the mechanism remains elusive, our results suggest that either ZMP effect on locomotion is developmental and occurs prior to the L4 stage, or ZMP accumulation from external AICAR does not affect the same tissues as endogenous ZMP build up in *atic-1(∆)*.

On a side note, because ZMP is an AMPK agonist [6], its riboside form AICAR is often added to culture media in different model organisms including *C. elegans* (*e.g.* [49, 50]), to trigger AMPK activation upon intracellular conversion to ZMP. Our results show that increase of ZMP is not the only metabolic consequence of AICAR treatment, therefore caution is recommended when attributing phenotypes to AMPK activation.

As to the locomotion defects in *adsl-1(∆)*, our results point to ADSL activity in the purine recycling pathway, given *ppat-1(∆)* does not suppress the locomotion phenotype, quite to the contrary. Furthermore, *adsl-1(∆)* is associated with significant damage in the structure of muscle myosin fibers, suggesting the locomotion defects are caused by muscle defects. While the underlying cause for these defects is yet to be determined, we hypothesize that the developmental delay in *adsl-1(∆)* is linked to muscle defects.

### Purine metabolic network in a metazoan model

The results we present show the functional conservation of both the *de novo* and recycling purine biosynthesis pathways in *C. elegans*. Furthermore, we characterized phenotypes associated with the deficiencies in the critical enzymes ADSL and ATIC. While several phenotypes are genetically separable (Figure 6), it stands out that the most severe phenotypes - lack of GSCs, developmental delay and muscle integrity - are associated with ADSL activity in the recycling pathway, whilst deficiencies in the *de novo* pathway are viable and lead to rather subtle phenotypes - as reduced fertility and slower locomotion. Noticeably, the deletion of *ppat-1* - which we show is required for the *de novo* pathway - produces no detectable phenotype in our culture conditions. The phenotypes observed in *atic-1(∆)*, locomotion defects and reduced fertility, are associated with ZMP accumulation, rather than to shortage in purine synthesis through the *de novo* pathway, as though the *de novo* pathway *per se* is dispensable. It might seem paradoxical that the purine *de novo* pathway is conserved while having no essential function. Indeed a number of biosynthesis pathways, namely amino-acids, have been lost during metazoan evolution [15] likely because amino-acids are available from the food source, as are nucleotides. However, these may not reflect the natural conditions in which *C. elegans* evolved; it is plausible that the *de novo* pathway is essential in different environments where purines are scarce. Actually, if one draws a parallel with yeast, it’s the severity of the *adsl-1(∆)* phenotypes that seems surprising, considering *Ade13* null mutants in *S. cerevisiae* are viable when cultured with adenine [51]. Given purines are available from the food source, one would have expected the worm to synthesize AMP and GMP despite ADSL deficiency. Indeed, external adenine and adenosine can be converted in AMP by Adenine phosphoribosyltransferase and Adenosine kinase respectively, both of which have orthologues in *C. elegans* (Supplementary Figure 1A), and AMP can be converted in GMP(Figure 1A). The severity of *adsl-1(∆)* phenotypes, suggests that ADSL became essential even when external purines are available. Importantly, in humans, purines are also available from the food source, and all but one diseases affecting purine metabolism are linked to enzymes in the recycling pathway; with the notable exception being AICA-ribosiduria - deficiency in ATIC, in which ZMP buildup may be a crucial disease factor [2, 3, 13]. The emerging picture is that in metazoans the less energetically costly recycling pathway evolved to become the predominant form of purine metabolism, at least in favorable environmental conditions. Hence, specific functions evolved to strongly rely on the recycling pathway, independently of the abundance of external purine sources. The defects associated with ADSL deficiency, as well as other recycling pathway deficiencies, could be the consequence of unbalanced wiring of the pathway, such as deregulation caused by intermediate metabolites, as much as insufficient synthesis of the final products AMP and GMP.

## Materials and Methods

### Caenorhabditis elegans strains and culture

Nematodes were maintained under standard culture conditions [52, 53], on nematode growth medium (NGM), with *Escherichia Coli* (strain OP50) as food source, kept at 20°C (except strains carrying the transgene ruIs32 [pie-1p::GFP::H2B + unc-119(+)], which were kept at 25°C).

The following *Caenorhabditis elegans* strains were used in the study: N2 (Bristol, wild-type reference strain) obtained from the Caenorhabditis Genetics Center (CGC), AZ212 *ruIs32 [pie-1p::GFP::H2B + unc-119(+)]* III, CF1903 *glp-1(e2141)* III, PE254 *feIs4 [sur-5p::luciferase::GFP + rol-6(su1006)]* V, VC2999 *R151.2(gk3067)* III*/hT2 [bli-4(e937) let-?(q782) qIs48]* (I,III) - parental strain for the hT2 balancer, WBX52 *adsl-1(tm3328)* I*/hT2 [bli-4(e937) let-?(q782) qIs48]* (I,III), WBX53 *adsl-1(tm3328)* I*; ppat-1(tm6344)* III*/hT2 [bli-4(e937) let-?(q782) qIs48] ppat-1(tm6344)* (I,III) (from WBX119 and WBX52), WBX116 *ppat-1(tm6344)* III, WBX119 *ppat-1(tm6344)* III, WBX117 *atic-1(tm6374)* IV, *atic-1(tm6374)* IV (from WBX119 and WBX117); WBX124 *atic-1(tm6374)* IV*; feIs4 [sur-5p::luciferase::GFP + rol-6(su1006)]* V (from PE254 and WBX116), WBX127 *ppat-1(tm6344)* I*; adsl-1(tm3328)* III*/hT2 [bli-4(e937) let-?(q782) qIs48] ppat-1(tm6344)* (I,III)*; feIs4 [sur-5p::luciferase::GFP + rol-6(su1006)]* V (from PE254 and WBX53), WBX125 *ppat-1(tm6344)* III*; atic-1(tm6374)* IV*; feIs4 [sur-5p::luciferase::GFP + rol-6(su1006)]* V (from PE254 and WBX116), WBX130 *adsl-1(tm3328)* I*/hT2 [bli-4(e937) let-?(q782) qIs48]* (I,III)*; feIs4 [sur-5p::luciferase::GFP + rol-6(su1006)]* V (PE254 and WBX53), WBX131 *ppat-1(tm6344)* III*; feIs4 [sur-5p::luciferase::GFP + rol-6(su1006)]* V (from PE254 and WBX116), WBX147 *ppat-1(tm6344) ruIs32 [pie-1p::GFP::H2B + unc-119(+)]* III (from AZ212 and WBX119), WBX149 *adsl-1(tm3328)* I*; ruIs32 [pie-1p::GFP::H2B + unc-119(+)]* III*/hT2 [bli-4(e937) let-?(q782) qIs48]* (I,III) (from AZ212 and WBX52) and WBX152 *adsl-1(tm3328)* I*; ppat-1(tm6344) ruIs32 [pie-1p::GFP::H2B + unc-119(+)]* III*/hT2 [bli-4(e937) let-?(q782) qIs48]* (I,III) (from WBX147 and WBX149). The *adsl-1(tm3328)*, *ppat-1(tm6344)* and *atic-1(tm6374)* deletion alleles - referred to as *adsl-1(∆)*, *ppat-1(∆)* and *atic-1(∆)* throughout the manuscript - were generated by the lab of Dr. Shohei Mitani, National Bioresource Project, Japan (NBRP), all strains carrying these alleles used in this study were outcrossed a minimum of four times.

Nematode synchronization: Adult worms were collected with M9 buffer in 15 ml conical tubes, centrifuged 1 min at 1000 rpm and supernatant was discarded. Worm bleach solution (NaOH 1 M, NaOCl 0.25 M) was added until carcasses dissolve and eggs release (~8 min). Eggs were washed three times with M9 buffer transferred onto NGM seeded plates.

RNAi: In order to generate the *adsl-1(RNAi)* feeding vector a PCR fragment was amplified w i t h o l i g o m e r s 5’ - C T T C T T C C G G C A G A AT C AT C C A G T G - 3’ a n d 5’ - TGACTGTCGGACGTACTCATTACC-3’ and inserted onto L4440 plasmid [54]. For RNAi experiments, HT115(DE3) bacteria transformed with either the *adsl-1(RNAi)* feeding vector, or L4440 (Control), were cultured overnight at 37°C in LB 150 mg/L Ampicillin and induced with 2 mM IPTG 1-2 h prior seeding onto NGM 2 mM IPTG 150 mg/L Ampicillin, in which synchronous populations of worms were cultured from hatching to adulthood.

### Yeast heterologue rescue assay

Yeast expression vectors were constructed by inserting full length cDNAs of *ppat-1* (amplified using oligomers 5’ - CACTAAATTACCGGATCAATTCGGGGGATCCATGTGTGGTATATTTGGGATTG-3’ and 5’- CATAACTAATTACATGATGCGGCCCTCCTGCAGTTATAAATCGATAGCAACTGG-3’), *adsl-1* (5’ - C G G G AT C C G TAT C AT G G C C T C C G A A G A C A A G T-3’ a n d 5’ - CCAATGCATGAATCAAACATCCAGCTGAACATTTCC-3’) and *atic-1b*(5’ - G A A G A T C T C G A C G A A T T G T G T G A A A T G A C C G A C - 3’ a n d 5’ - ATAAGAATGCGGCCGCATCTAATGGTGGAAGAGACGAAGTC-3’) onto pCM189 plasmid[55]. *ade4∆* yeast strain, genotype *ade4::KanMX4; his3∆1; leu2∆0; lys2∆0; ura3∆0*, was used to test heterologous rescue by *ppat-1*, *ade13∆* yeast strain, genotype *ade13::kanMX4; his3∆; leu2∆; ura3∆*, was used to test heterologous rescue by *adsl-1* and *ade16∆ ade17∆* double mutant, genotype *ade16::KanMX4; ade17::KanMX4; his3∆1; leu2∆0; ura3∆0; lys2∆0* was used to test heterologous rescue with *atic-1b*.

### Imaging

For adult morphology, DIC (Differential Interference Contrast, *a.k.a.* Nomarski) images were taken with a Leitz Diaplan microscope. Adult worms were mounted in M9 buffer, on 2% agarose cushion. Images were acquired with VisiCam 5.0 camera, analyzed and processed with Fiji software. For imaging of *pie-1p::GFP::H2B* germline reporter, synchronized adult worms were mounted in M9 Buffer 5 mM Levamisole onto 2% agarose cushion. Images were acquired with an Olympus IX81 microscope under an 40X oil-immersion objective, with Hamamatsu Orca R2 camera. Images were analyzed with Fiji software. For size measurements synchronized populations of young adult worms were used, each worm was measured 3 times on three different pictures. Images were acquired on a Nikon SMZ18 stereo microscope with VisiCam 5.0 camera, analyzed and processed with Fiji software.

### Immunostainings

Immunostaining was performed on synchronized populations of worms, N2 and *de novo* pathway mutants were fixed 72 h after synchronization, *adsl-1(∆)* and *ppat-1(∆); adsl-1(∆)* were fixed 96h after synchronization. Fixation protocol was adapted from Benian et al. [56]. Worms were collected in ice cold PBS buffer on 1.5 ml tubes, centrifuged and supernatant was discarded, and fixation solution was added (made fresh, 1ml of MRBW buffer KCl 320 mM, NaCl 80 mM, EGTA 40 mM, Spermidine 20 mM, Pipes 60 mM was added to 2.8 ml of Methanol and 200 µl of Formaldehyde 20%), followed by five cycles of freezing in liquid N2 and thawing and 1 h incubation at 4°C with gentle agitation, fixation solution was replaced by a permeabilization solution (PBS, Triton X-100 2%, ß-mercapto-ethanol 1%) and incubated 1h at room temperature, permeabilization solution was renewed and incubated 1h at room temperature, followed by three washes in PBSt (PBS, Triton X-100 0.5%), after which worms were incubated with primary antibody (diluted 1:1000 in PBSt 5% Fetal Calf Serum) overnight at 4°C with gentle agitation, washed three times in PBSt and incubated with secondary antibody (diluted 1:500 in PBSt 5%FCS). Monoclonal antibody 5-6 (Developmental Studies Hybridoma Bank) directed against muscle myosin was used as primary antibody, anti-mouse Alexa555(Molecular Probes) was used as secondary antibody, worms were mounted on Fluoroshield (Sigma). Images were acquired with a Leica DM6000 TCS SP5 confocal microscope using LAS AF (Leica) software.

### Metabolic profiling by HPLC

For an extraction, ~7500 young adult worms, 72 h after synchronization, were collected with ice cold water, onto 15 ml conical tubes, and centrifuged at 1000 rpm, 1 min and 4°C, supernatant was discarded eliminating most bacteria. The nematode pellet was washed with 10 ml of cold Hepes 10 mM and centrifuged. The worm pellet was resuspended in Hepes 10 mM, 1 ml of worm suspension is transferred onto 4 ml of absolute ethanol at 80°C, incubated for 5 min and cooled on ice for 10 min. Samples were desiccated on a rotovapor apparatus, the dry residue was resuspended in 350 µl of HyClone Molecular Biology-Grade Water (GE Healthcare) and centrifuged at 21000 g, 5 min at 4°C, supernatant was transferred and centrifuged again at 21000 g, 1 h at 4°C. The last supernatant was used for HPLC. Metabolites were separated by ionic chromatography on an ICS3000 chromatography station (Dionex, Sunnyvale, CA), and measured by an UV diode array (Ultimate 3000 RS, Dionex). Metabolites separation was done on a carbopacPA1 column (250 × 2 mm; Dionex) using a sodium acetate gradient [57]. Metabolite levels were normalized relative to dry weight; the same amount of worms used in the extractions were collected in ice cold water, 72h after synchronization, centrifuged, washed 5 times with water, allowed to evaporate for 4 days at 70°C and weighed.

### Lifespan

Upon synchronization worms were transferred onto a new plate every 24 h, and scored. Immobile worms unresponsive to touch were considered dead.

### Locomotion and Behavioral assays

Locomotion speed measurement was adapted from [29]; images were acquired on a Leica Z16 APO Macroscope and processed with Fiji software, worm displacement was measured by tracking the pharynx, one frame per second.

Aldicarb (Sigma) and levamisole (Sigma) assays were performed as described in [34], synchronized adult worms were placed on 1 mM Aldicarb or 0.4 mM Levamisole plates for 2 h, after which plates were tapped 10 times and worm movement was scored.

Mechanosensory response assay was performed as described in [29], synchronized adult worms were isolated on plates without food for 3 min, and stimulated with an eyelash on the posterior pharynx, and scored for avoidance behavior, each worm was stimulated 10 time 3 min apart.

### Post embryonic development

Monitoring of post-embryonic development was performed as described by Olmedo *et al.* [25]. In brief, upon synchronization, 20 embryos per µl in M9 buffer overnight with agitation, allowing L1 larvae to hatch. A single L1 larvae was added per well in 96-well plates with S-Basal media, 10 g/L E. coli OP50 (wet weight) and 100 µM D-Luciferin. The luminescence was measured with a Berthold Centro LB960 XS3 during 1 second at 5-min intervals, within a temperature controlled Panasonic MIR-154 incubator.

### Statistics

Statistical analysis was done using Graphpad, Excel, Numbers and R softwares. We consider statistical test to be significant when pValue <0.01.

## Acknowledgements

Some *C. elegans* strains were provided by the CGC (https://cgc.umn.edu), which is funded by NIH Office of Research Infrastructure Programs (P40 OD010440). We thank Wormbase, which is supported by National Institutes of Health (NIH) grant U41 HG002223. This work was supported by an AFM-Téléthon Trampoline grant (ref. 18313). RM is supported by AFM-Téléthon PhD scholarship (ref. 20450). RM stay in M. A-S. lab supported by a Short Term Scientific Mission financed by GENiE (COST Action BM1408). M.A-S and M.O are supported by the Ramón y Cajal program of the Spanish Ministerio de Economía y Competitividad, RYC-2010-06167 and RYC-2014-15551, respectively. Confocal microscopy, and locomotion assay imaging were done in the Bordeaux Imaging Center, a service unit of the CNRS-INSERM and Bordeaux University, member of the national infrastructure France BioImaging. Deletion alleles *adsl-1(tm3328)*, *ppat-1(tm6344)* and *atic-1(tm6374)* were generated by the National Bioresource Project, Japan, directed by Shohei Mitani. We thank Yuji Kohara for providing cDNAs. We thank the members of the B. D-F. lab for helpful discussions, and particularly M. Moenner for critically reading the manuscript.

---

**Supplementary Figure 1.**
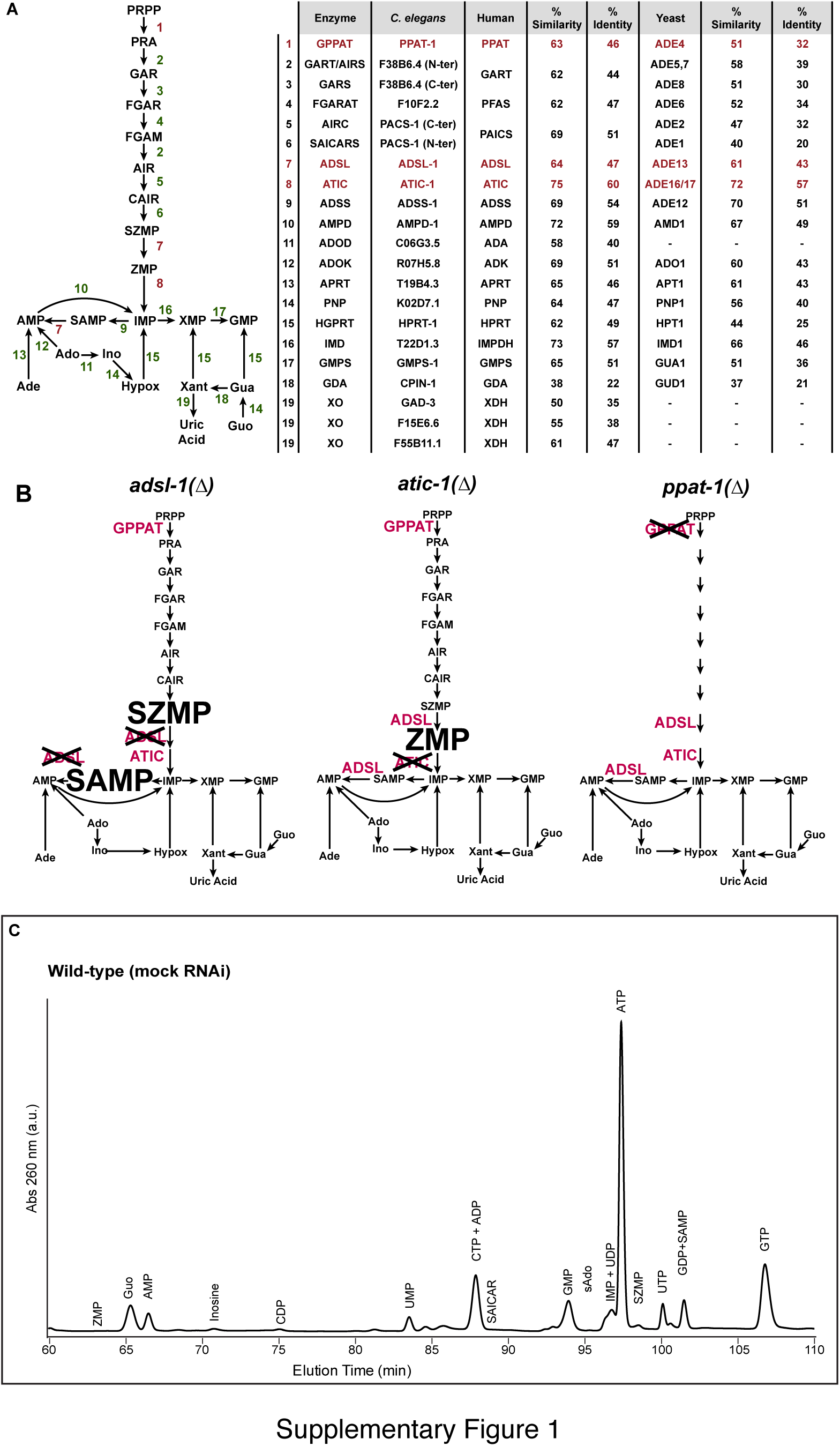
Purine biosynthesis pathway conservation in *C. elegans*. (A) Schematics of the Purine Biosynthesis pathways in *C. elegans* based on sequence homology, and table of comparison with both human and yeast (*S. cerevisiae*) orthologues. Enzymes subject to functional analysis are in red. (B) Schematics of predicted purine biosynthesis pathways in *adsl-1(∆)*, *atic-1(∆)* and *ppat-1(∆)* mutants; large font represents intermediate metabolite accumulation. (C) Representative example of a chromatogram, absorbance at 260 nm, of wild-type worms, in mock (control) RNAi condition.

**Supplementary Figure 2.**
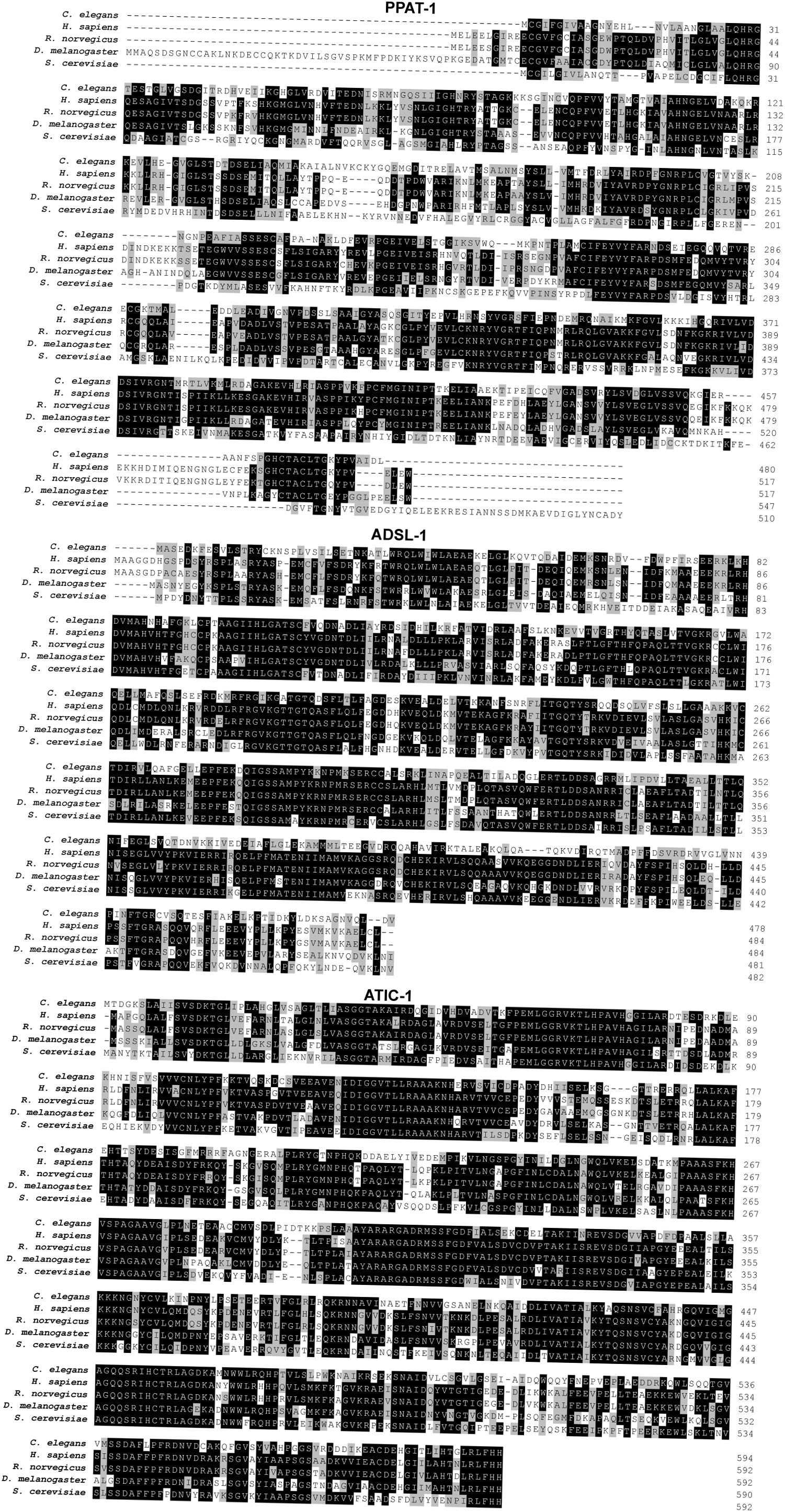
Sequence alignment of *C. elegans* PPAT-1, ADSL-1 and ATIC-1 with closest orthologue in humans, rat (*Rattus norvegicus*), fly (*Drosophila melanogaster*) and yeast (*S. cerevisiae*). Grey boxes indicate similar residues, black boxes indicate identical residues.

**Supplementary Figure 3.**
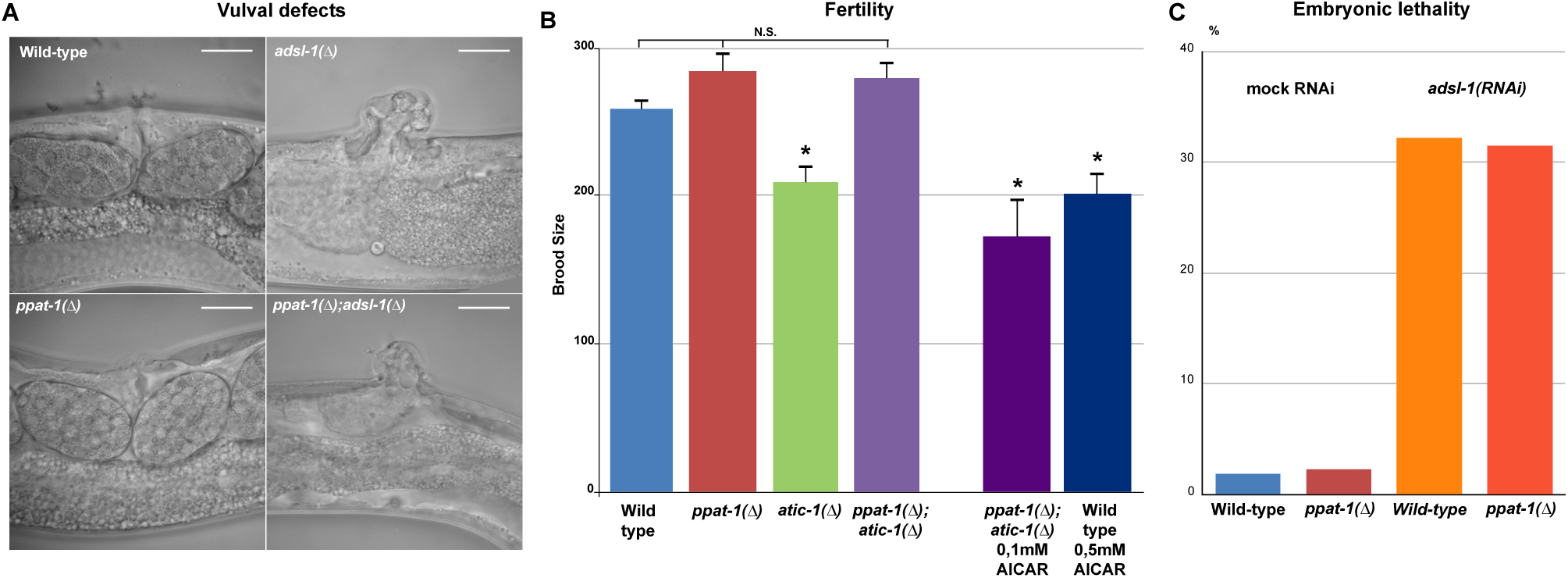
Developmental defects in purine mutants. (A) Representative DIC micrographs of the vulva region in young adults of wild-type, *ppat-1(∆)*, *adsl-1(∆)* and *ppat-1(∆); adsl-1(∆)* genotypes (bar= 20µm). (B) Graph depicting the average number of eggs laid by the *de novo* pathway mutants, and AICAR treated *ppat-1(∆); atic-1(∆)* (0.1 mM) and wild-type (0.5 mM) worms (error bar - standard error of the mean; n= 30 in all samples; * - statistically significant difference relative to wild-type - Student’s t-test). (C) Graph displaying the percentage of embryonic lethality (unhatched eggs) upon *adsl-1(RNAi)* and mock (control) RNAi in both wild-type and *ppat-1(∆)*.

**Supplementary Figure 4.**
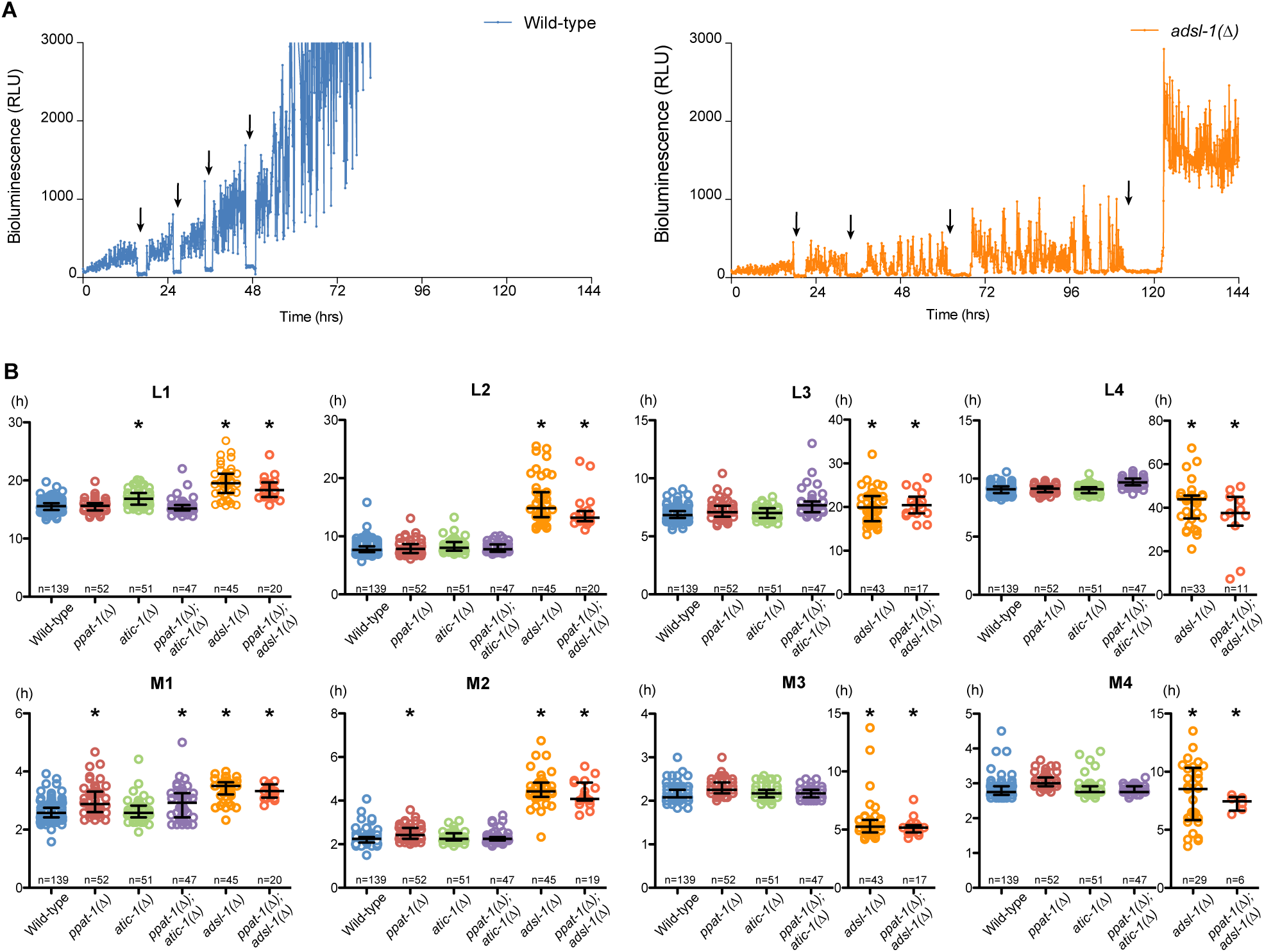
(A) Representative bioluminescence profiles of a single wild-type (left, blue line) and *adsl-1(∆)* mutant (right, orange line) individuals over time. Arrows indicate drop in luminescence observed during lethargus phase. (B) Detailed representation of the primary data presented in figure 3A, each circle corresponds to an individual observation, bars indicate median, 2nd and 3rd quartiles. L1 through L4 refer to the four larval stages and M1 though M4 refer to the lethargus phases following each larval stage. For better visualization *adsl-1(∆)* and *ppat-1(∆); adsl-1(∆)* in L3, L4, M3 and M4 are represented on a different time scale. (* - statistically significant difference compared to wild-type; one-way ANOVA).

**Supplementary Figure 5.**
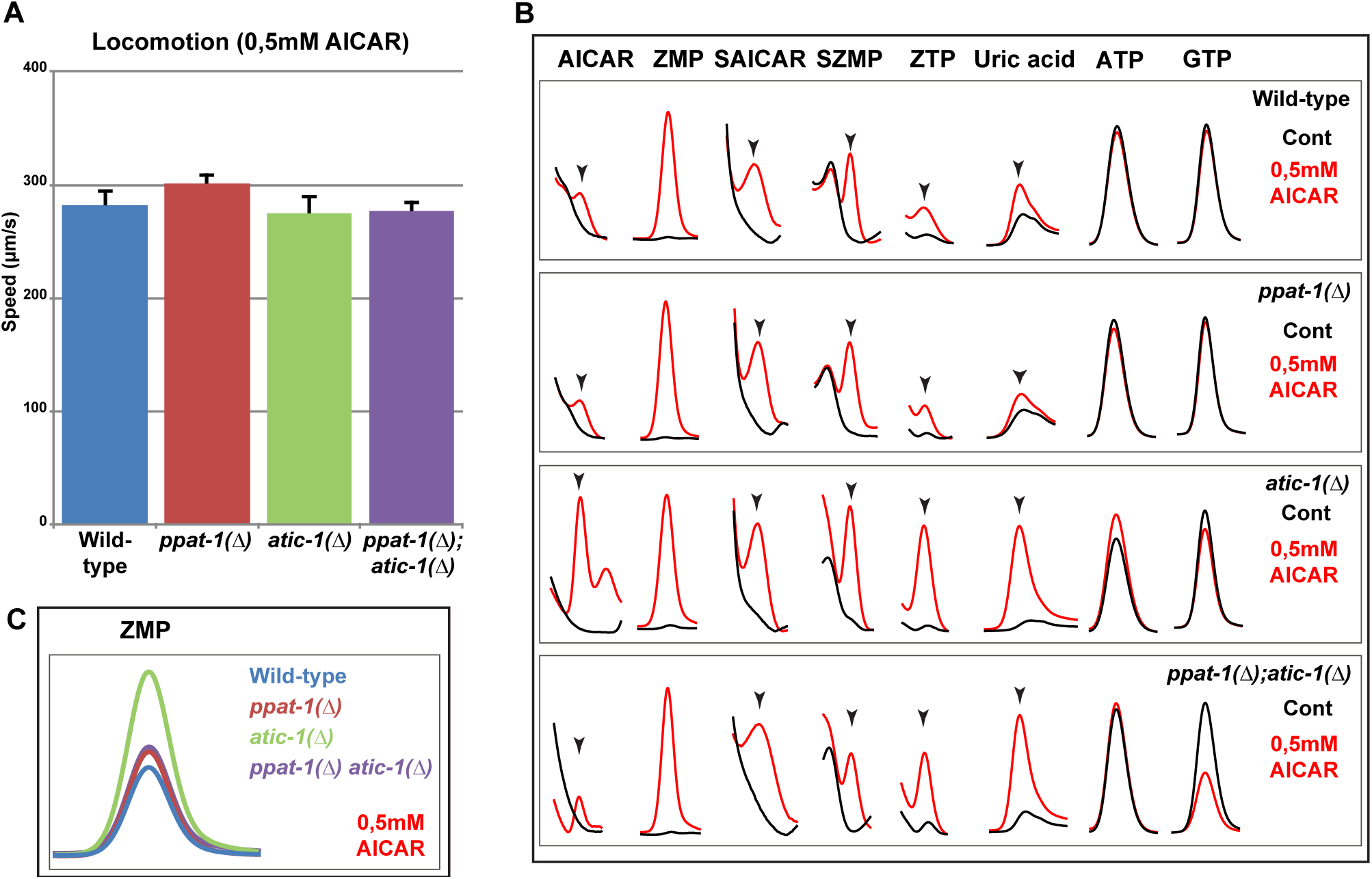
AICAR treatment and locomotion. (A) Graph depicting the average locomotion speed in *de novo* pathway mutants treated with AICAR (error bar - standard error of the mean; n= 30 in *ppat-1(∆)* and *atic-1(∆),* n= 40 in wild-type and *ppat-1(∆); atic-1(∆)*; * - statistically significant difference relative to wild-type - Student’s t-test). (B) Zoom in on HPLC chromatogram peaks of specific metabolites AICAR, ZMP, SAICAR, SZMP, ZTP, uric acid, ATP and GTP, upon 0.5 mM AICAR treatment. (C) Zoom in on HPLC chromatogram peaks of ZMP upon 0.5 mM AICAR treatment, with the four genotypes analyzed represented on the sale scale.

**Supplementary Figure 6.**
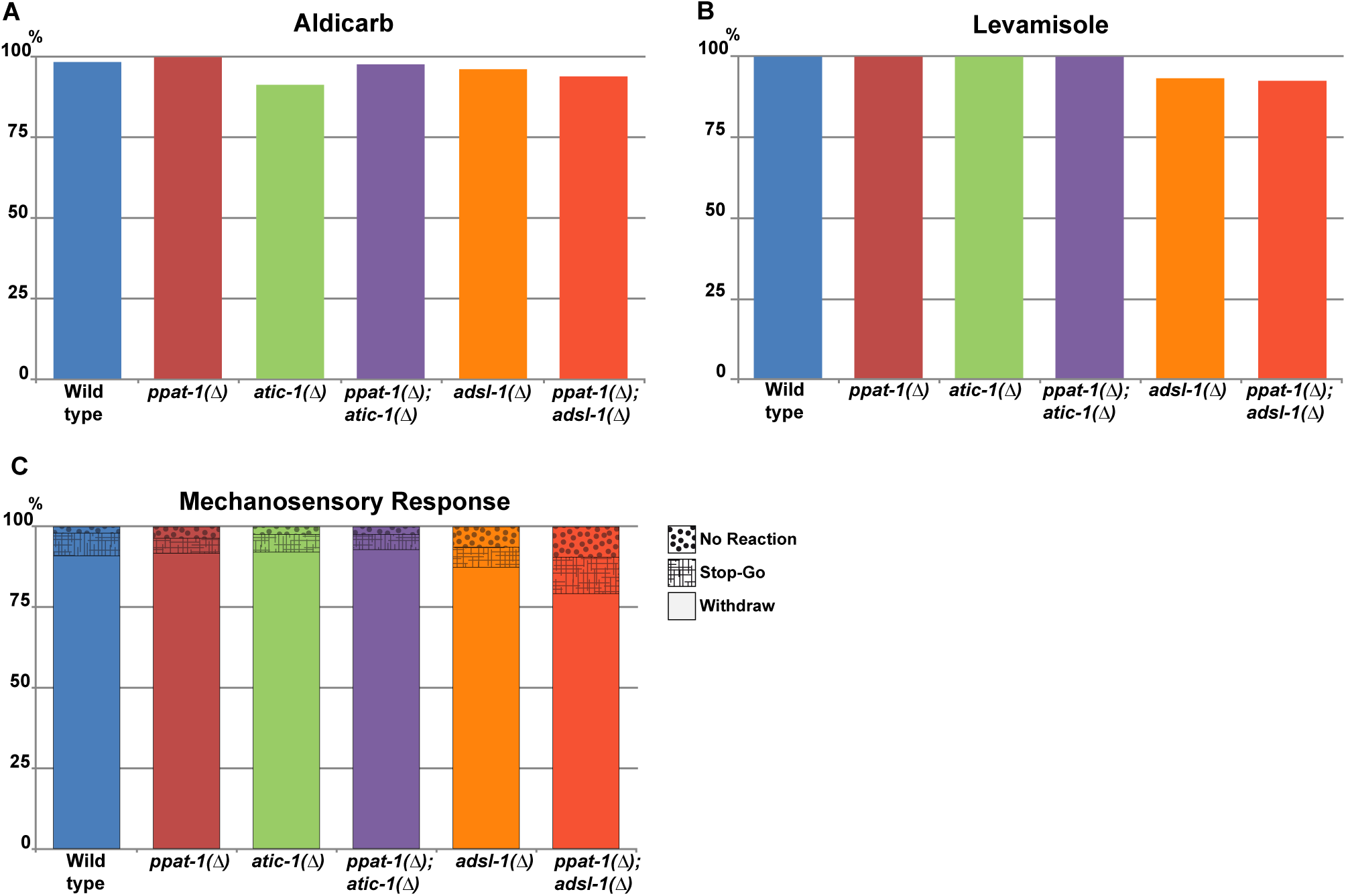
Neuromuscular defects in purine mutants. (A) Percentage of paralysed worms upon Aldicarb treatment, in all studied genotypes. (B) Percentage of paralysed worms upon Levamisole treatment, in all studied genotypes. (C) Graph presenting the percentage of observed responses to mechanosensorial stimulation in all studied genotypes.

